# Brevicidine, a bacterial non-ribosomally produced cyclic antimicrobial lipopeptide with a unique *modus operandi*

**DOI:** 10.1101/2022.10.07.511251

**Authors:** Xinghong Zhao, Xinyi Zhong, Hongping Wan, Lu Liu, Xu Song, Yuanfeng Zou, Lixia Li, Renyong Jia, Juchun Lin, Huaqiao Tang, Gang Ye, Jianqing Yang, Shan Zhao, Yifei Lang, Zhongqiong Yin, Oscar P. Kuipers

## Abstract

Due to the accelerated appearance of antibiotic-resistant (AMR) pathogens in clinical infections, new first-in-class antibiotics, operating via novel modes of action, are desperately-needed. Brevicidine, a bacterial non-ribosomally produced cyclic lipopeptide, has shown potent and selective antimicrobial activity against Gram-negative pathogens. However, before our investigations, little was known about how brevicidine exerts its potent bactericidal effect against Gram-negative pathogens. In this study, we find that brevicidine has potent antimicrobial activity against AMR *Enterobacteriaceae* pathogens, with a MIC value ranging between 0.5μM (0.8mg/L) and 2μM (3.0mg/L). In addition, brevicidine showed potent anti-biofilm activity against the *Enterobacteriaceae* pathogens, with same 100% inhibition and 100% eradication concentration of 4μM (6.1mg/L). Further mechanistic studies showed that brevicidine exerts its potent bactericidal activity via interacting with lipopolysaccharide in the outer membrane, targeting phosphatidylglycerol and cardiolipin in the inner membrane, and dissipating the proton motive force of bacteria. This results in metabolic perturbation, including inhibition of adenosine triphosphate synthesis, inhibits the dehydrogenation of nicotinamide adenine dinucleotides, accumulation of reactive oxygen species in bacteria, and inhibition of protein synthesis. Lastly, brevicidine showed a good therapeutic effect in a mouse peritonitis–sepsis model. Our findings pave the way for further research on clinical applications of brevicidine, to combat the prevalent infections caused by AMR Gram-negative pathogens worldwide.

## Introduction

Due to the accelerated appearance of antimicrobial resistance (AMR) in bacterial pathogens, many bacterial infections have become increasingly difficult, or even impossible, to treat with conventional antimicrobials. The O’Neill report on AMR, published in 2014, predicted that deaths attributable to resistant infections will reach 10 million annually by 2050 (1, 2). In addition, due to the COVID-19 pandemic, the AMR situation is getting worse since increasing evidence has emerged that the COVID-19 pandemic also contributes to AMR (2–6). Despite the AMR situation being so urgent, the number of newly approved antimicrobials has been steadily decreasing over the past 20 years, especially those for treating Gram-negative pathogen infections (7–9). Bacteria produce a wealth of non-ribosomally produced peptide (NRP) antimicrobials, including clinically approved antimicrobial cyclic lipopeptides, such as vancomycin, daptomycin, and colistin (10). This shows that the group of cyclic lipopeptides makes up a rich source of molecules able to control pathogenic bacterial infections.

Brevicidine (Bre), a bacterial non-ribosomally produced cyclic lipopeptide, was first isolated from the *Brevibacillus laterosporus* DSM25 in 2018 (11). This cyclic lipopeptide contains 12 amino acids (4-Methyl-Hexanoyl-D-Asn-D-Tyr-D-Trp-D-Orn-Orn-Gly-D-Orn-Trp-Thr-Ile-Gly-Ser) with a 4-Methyl-Hexanoyl chain at its N-terminus and a lactone bond between Thr9 and Ser12 (Fig. 1) (11, 12). In addition, Nathaniel et al reported the total synthetic route of brevicidine in 2022 (13), which makes it more attractive for antibiotic development. Brevicidine displays potent and selective antimicrobial activity against Gram-negative pathogens, including *Enterobacter cloacae, Escherichia coli, Pseudomonas aeruginosa*, and *Klebsiella pneumoniae* (11, 14), which are listed as “critical” priority pathogens that need R&D of new antimicrobials by the World Health Organization (WHO) (15). Notably, brevicidine showed no cytotoxicity and hemolytic activity at a relatively high concentration of 128μg/ml (11, 14), indicating brevicidine is a promising antimicrobial candidate. In our previous study, we found that brevicidine dissipated the proton motive force of bacteria. However, before the present investigations, little was known about how brevicidine exerts its potent bactericidal effect against Gram-negative pathogens. In addition, the antimicrobial activity against AMR pathogens and the therapeutic effect *in vivo* of brevicidine still needs to be investigated before it is considered for a clinical trial.

**Fig. 1.**
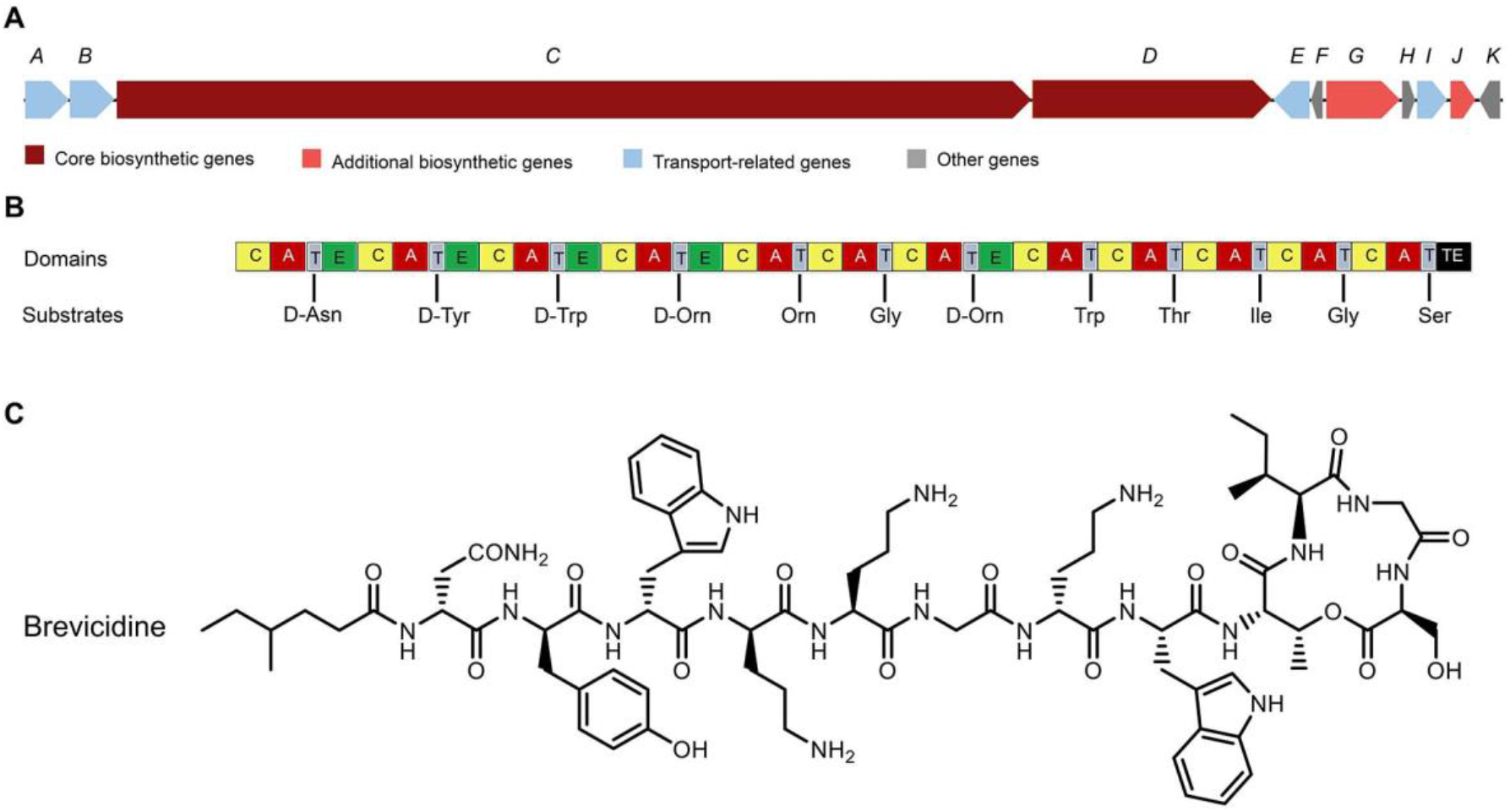
The structure of brevicidine and the predicted biosynthetic gene cluster (14). **A**, The non-ribosomal peptide synthetases genes harbored by the *Brevibacillus laterosporus* DSM 25 genome; **B**, the catalytic domains encoded by the gene cluster, and the substrates incorporated by the respective modules. Domains: A, adenylation; T, thiolation; C, condensation; E, epimerization; TE, thioesterase; **C**, the structure of brevicidine.

In this study, we aimed to identify the precise molecular mechanism of action by which brevicidine kills Gram-negative pathogens. Moreover, the anti-biofilm activity against *E. coli*, the antimicrobial activity against AMR *Enterobacteriaceae*, and the *in vivo* therapeutic effect of brevicidine were also investigated. The results showed that brevicidine exerts its potent bactericidal effect by interacting with lipopolysaccharide (LPS) in the outer membrane as well as targeting phosphatidylglycerol (PG) and cardiolipin (CL) in the inner membrane, and subsequently disrupting the proton motive force of Gram-negative pathogens. This leads to adenosine triphosphate (ATP) synthesis inhibition, nicotinamide adenine dinucleotide (NADH) and reactive oxygen species (ROS) accumulation, and bacteria death. In addition, brevicidine showed potent antimicrobial activity against AMR Gram-negative pathogens and good anti-biofilm activity against Gram-negative pathogen. Notably, brevicidine showed a good therapeutic effect in an *E. coli*-induced mouse peritonitis–sepsis model. These results will pave the way for further research on the clinical applications of brevicidine.

## Results and Discussion

### Brevicidine shows potent antimicrobial activity against antibiotic-resistant (AMR) *Enterobacteriaceae* pathogens

The antimicrobial activity of brevicidine against AMR *Enterobacteriaceae* pathogens was measured by a MIC assay. The results show that brevicidine has potent antimicrobial activity, with a MIC value of 0.5μM (0.8mg/L) to 2μM (3.0mg/L) (Table 1), against *Enterobacteriaceae* pathogens, including the tested colistin-resistant strains and multidrug-resistant strains (16). These results indicate that brevicidine has no cross-resistance with colistin, ampicilin, ceftriaxone, cefotaxime, aztreonam, gentamicin, ofloxacin, amoxicillin, streptomycin, lincomycin, doxycycline, or ciprofloxacin, which makes it a more attractive antimicrobial candidate for treating AMR Gram-negative pathogen infections.

**Table 1.** MIC values of brevicidine against *Enterobacteriacea*e bacterial pathogens.

| Microorganism | MIC [μM (mg/L)] | | | |
| --- | --- | --- | --- | --- |
|  | Brevicidine | Colistin | Amikacin | Ampicilin |
| *Escherichia coli* ATCC25922 | 0.5 (0.8) | 2 (2.3) | 2 (1.2) | 8 (2.8) |
| *Escherichia coli* CVCC3749 | 1 (1.5) | 2 (2.3) | 2 (1.2) | > 256 (89) |
| *Escherichia coli* B2<sup>a</sup> | 1 (1.5) | 8 (9.2) | 32 (19) | > 256 (89) |
| *Escherichia coli* 16QD<sup>a</sup> | 1 (1.5) | 8 (9.2) | 8 (4.7) | > 256 (89) |
| *Escherichia coli* EPZC17-45<sup>b</sup> | 1 (1.5) | 2 (2.3) | 4 (2.3) | > 256 (89) |
| *Escherichia coli* EPZC17-22<sup>b</sup> | 1 (1.5) | 1 (1.6) | 2 (1.2) | > 256 (89) |
| *Escherichia coli* E21-5<sup>b</sup> | 0.5 (0.8) | 2 (2.3) | 2 (1.2) | > 256 (89) |
| *Escherichia coli* E21-71<sup>b</sup> | 0.5 (0.8) | 4 (4.6) | 4 (2.3) | > 256 (89) |
| *Escherichia coli* E21-75<sup>b</sup> | 0.5 (0.8) | 2 (2.3) | 8 (4.7) | > 256 (89) |
| *Escherichia coli* E21-107<sup>b</sup> | 0.5 (0.8) | 1 (1.6) | 4 (2.3) | > 256 (89) |
| *Enterobacter cloacae* LMG02783<sup>c</sup> | 1 (1.5) | > 64 (74) | 8 (4.7) | > 256 (89) |
| *Klebsiella pneumoniae* LMG20218 | 2 (3.0) | 2 (2.3) | 4 (2.3) | > 256 (89) |
| *Staphylococcus aureus* ATCC 43300<sup>d</sup> | 32 (48.6) | 32 (37) | 4 (2.3) | 16 (5.6) |
| *Enterococcus faecium* LMG16003<sup>e</sup> | > 32 (48.6) | > 32 (37) | 256 (150) | 2 (0.7) |

<sup>a</sup>*Escherichia coli* carrying mcr-1, blaNDM-5, mdx genes, etc., which is resistant against colistin, carbapenems, rifampin, etc. <sup>b</sup> *Escherichia coli* (clinical isolation) carrying multidrug resistance genes, *Escherichia coli* EPZC17-45, resistant to ampicillin, ceftriaxone, cefotaxime, aztreonam, gentamicin, doxycycline, and ciprofloxacin; *Escherichia coli* EPZC17-22, resistant to ampicillin, ceftriaxone, cefotaxime, aztreonam, gentamicin, doxycycline, and ciprofloxacin; *Escherichia coli* E21-5, resistant to ampicillin, amoxicillin, ceftriaxone, gentamicin, cefotaxime, streptomycin, doxycycline; *Escherichia coli* E21-71, resistant to ampicillin, amoxicillin, ceftriaxone, cefotaxime, doxycycline, and ciprofloxacin, and ofloxacin; *Escherichia coli* E21-75, resistant to ampicillin, amoxicillin, ceftriaxone, cefotaxime, doxycycline, ofloxacin, and ciprofloxacin; *Escherichia coli* E21-107, resistant to ampicillin, amoxicillin, ceftriaxone, gentamicin, cefotaxime, streptomycin, doxycycline, and lincomycin. <sup>c</sup>*Enterobacter cloacae* carrying mcr-9 gene, which is a potent colistin-resistant gene. <sup>d</sup> *Staphylococcus aureus* resistant to oxacillin and methicillin. <sup>e</sup> *Enterococcus faecalis* resistant to vancomycin.

### Brevicidine shows potent anti-biofilm activity against *E. coli*

Bacterial biofilm formation could increase the resistance of pathogens to conventional antimicrobials, which could decrease the therapeutic effect of antimicrobials (17, 18). Therefore, the anti-biofilm activity of antimicrobials plays a vital role in their therapeutic effect on biofilm-formation bacterial pathogen infections. In this study, the anti-biofilm activity of brevicidine against *E. coli* was evaluated according to the protocol described previously (19). Colistin (Col) was used as an anti-biofilm antimicrobial control, while amikacin (Amk) was used as a control antimicrobial without anti-biofilm activity. The results show that brevicidine has potent anti-biofilm activity against *E. coli*, with the same 100% inhibition and eradication concentration of 4μM (6.1 mg/L), which is better than the anti-biofilm activity of colistin against *E. coli* (Fig. 2). These results demonstrate that brevicidine shows high potential for the development of therapeutic antimicrobial.

**Fig. 2.**
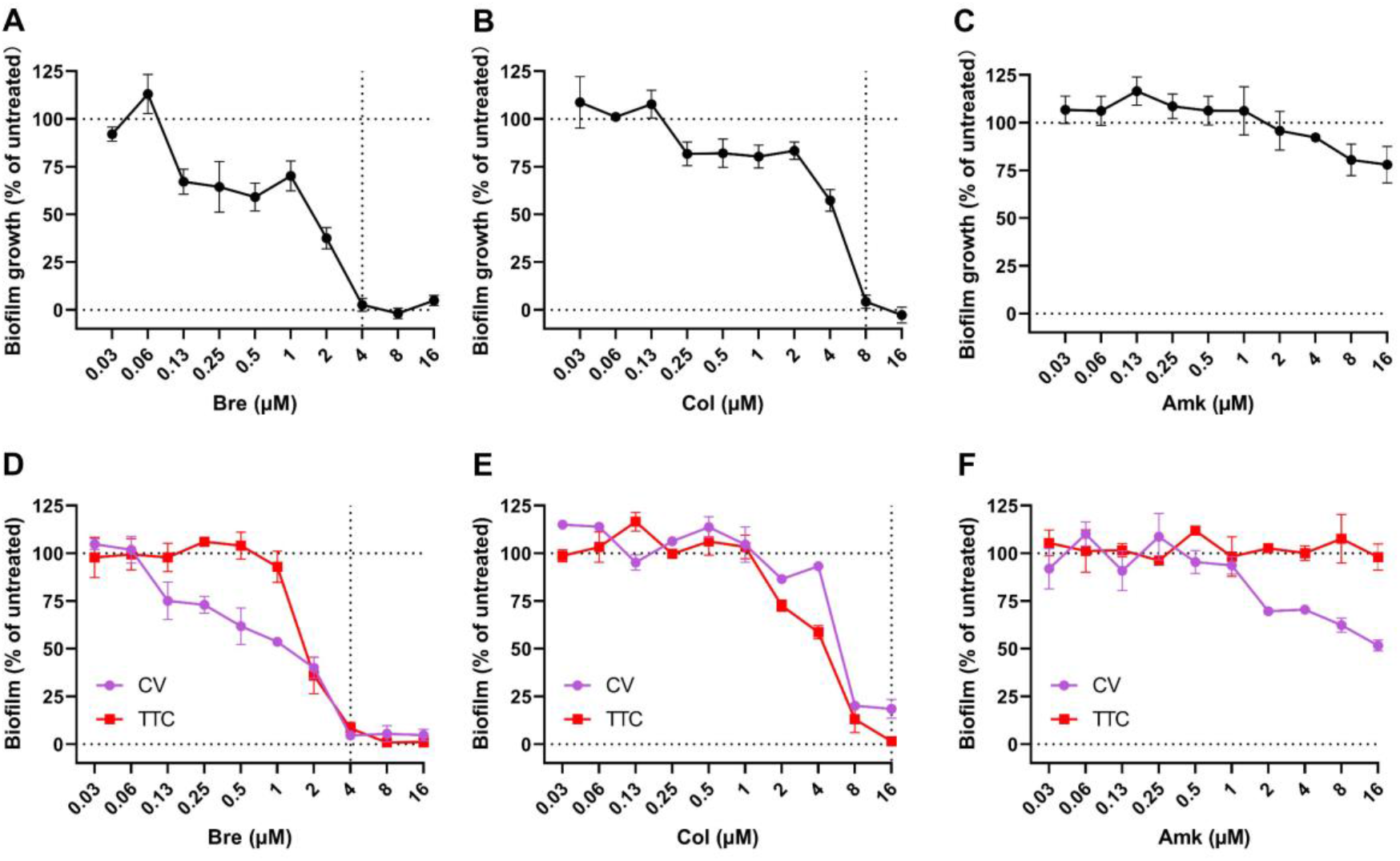
Brevicidine showed potent anti-biofilm activity against *E. coli* O101. **A**, Biofilm inhibition of various antimicrobials (brevicidine, colistin, and amikacin) versus *E. coli* O101. Biofilm inhibition was assessed by incubating the antimicrobials with bacteria in growth media for 24 h, discarding the growth media and staining the adhered biomass with CV. The percentage of biofilm inhibition is reported relative to untreated bacteria (defined as 100%) and sterility control wells (defined as 0%). All data were presented as means ± Standard Deviation. **B**, Biofilm eradication assay results for various antimicrobials (brevicidine, colistin, and amikacin) versus *E. coli* O101. Biofilm biomass, quantified based on CV staining, is indicated in pink. Biofilm metabolism, quantified based on TTC metabolism, is indicated in red. All data were presented as means ± Standard Deviation.

### Brevicidine disrupts the electron transport chain of Gram-negative bacteria

In previous studies, brevicidine had shown bacterial lytic activity at a concentration of 10×MIC (11, 14). However, it did not disrupt the membrane at a concentration of 2×MIC, evidenced by an atomic force microscopy assay. Therefore, the authors suggested that a more in-depth mechanism study is needed to characterize the targets as well as the membrane disruption effect of brevicidine (11). In our previous study, we found that brevicidine, at a concentration of 1×MIC, could dissipate the proton motive force of *E. coli*, evidence by a DiSC_3_(5) assay (14). However, before our investigations, little is known about the molecular mechanism of brevicidine on gram-negative bacteria. In this study, to investigate the killing capacity of brevicidine at concentrations of low MIC values, 4×MIC, 2×MIC, and 1×MIC, a time-killing assay was performed as the previously described method (20, 21). Brevicidine quickly decreased the population of *E. coli* at a concentration of 4×MIC, which killed all of the tested bacteria in 1h. Interestingly, the population of *E. coli* did not decrease under the treatment of brevicidine at concentrations of 2×MIC and 1×MIC within 30min (Fig. 3A). These results indicate that brevicidine has different modes of action between high and low MIC concentrations; it shows bacterial lytic activity at concentrations of 4-fold to 10-fold MIC, while it does not disrupt the membrane at concentrations of 1-fold to 2-fold MIC (11, 14). The atomic force microscopy, which was used in the previous study (11), is not a common technique for investigating the membrane effect of antimicrobials because of its low resolution. Therefore, in this study, we performed several fluorescent probe assays, scanning electron microscopy (SEM) assay, and transmission electron microscopy (TEM) assay to get insight into the effect of brevicidine, at low MIC level concentration, on the Gram-negative bacterial membrane.

**Fig. 3.**
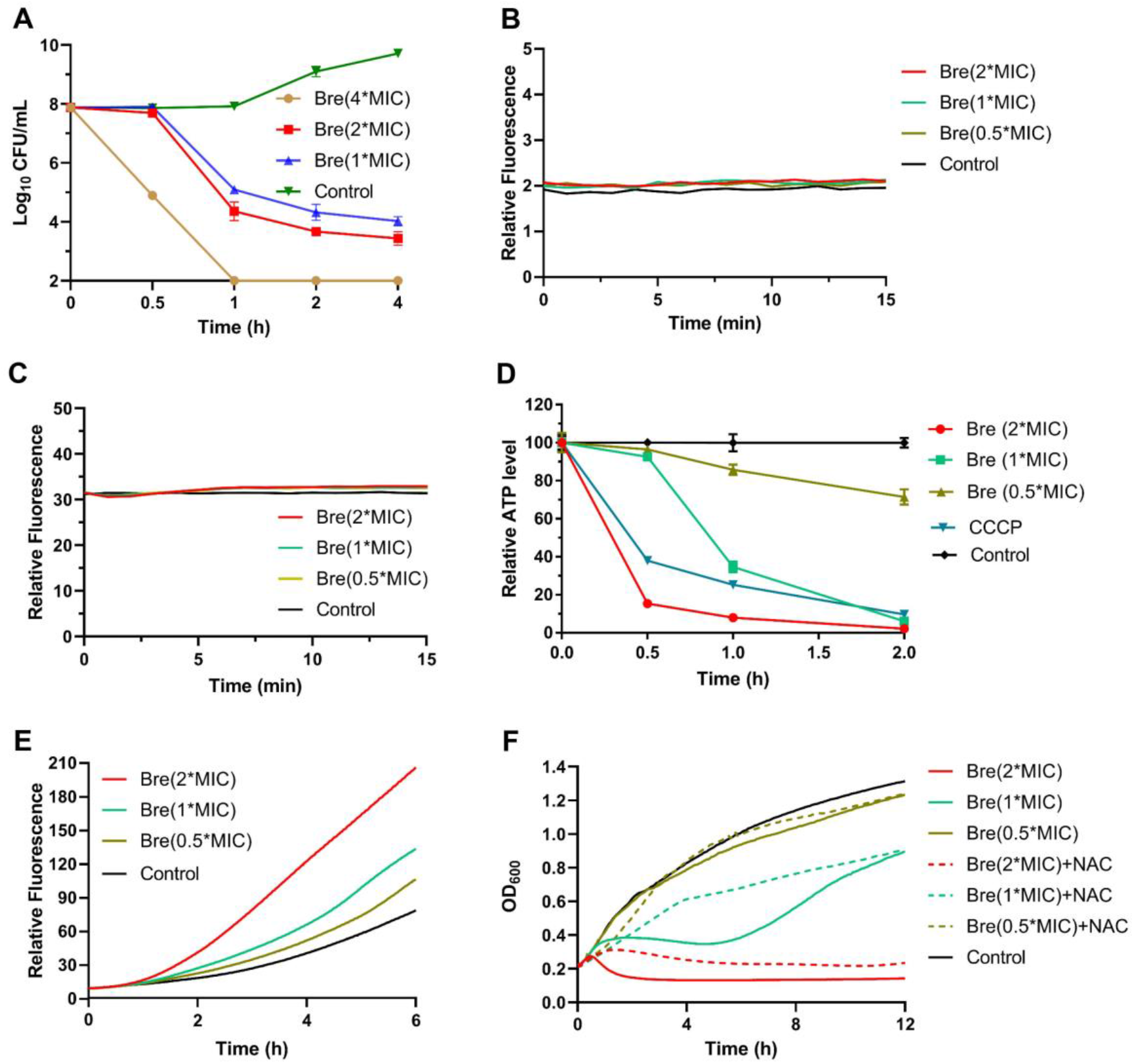
Brevicidine kills Gram-negative bacteria via dissipating their electron transport chain. **A**, time-killing curves of brevicidine at concentrations of 4×MIC, 2×MIC, and 1×MIC against *E. coli* O101. **B**, *E. coli* O101 cells pretreated with propidium iodide were exposed to antimicrobial peptides brevicidine at concentrations of 2×MIC, 1×MIC, and 0.5×MIC, and the extent of membrane leakage was visualized as an increase in fluorescence. **C**, NPN fluorescence in *E. coli* O101 cells upon exposure to brevicidine at concentrations of 2×MIC, 1×MIC, and 0.5×MIC. Representative examples from three technical replicates are shown. **D**, ATP concentration of *E. coli* O101 cells treated with brevicidine at concentrations of 2×MIC, 1×MIC, and 0.5×MIC. **E**, Total ROS accumulation in *E. coli* O101 cells treated with brevicidine at concentrations of 2×MIC, 1×MIC, and 0.5×MIC. Representative examples from three technical replicates are shown. **F**, Growth curves of *E. coli* O101 exposed to brevicidine at concentrations of 2×MIC, 1×MIC, and 0.5×MIC. Exogenous addition of NAC (6 μM) diminished the bactericidal activity of brevicidine.

The effect of brevicidine on the membrane integrity of Gram-negative bacteria was investigated by using the fluorescent probes N-Phenyl-1-naphthylamine (NPN) and propidium iodide (PI). NPN is an outer-membrane barrier permeant fluorescent probe (22). If the outer membrane is disrupted, the fluorescent probe can enter, reaching the phospholipid layer and resulting in a significant increase of fluorescence. The fluorescent probe PI is permeant to membrane intact bacteria, and the disruption of membrane integrity can lead to a prominent increase in fluorescence (20). There was no fluorescence increase observed for both brevicidine-treated NPN-containing cell suspensions and brevicidine-treated PI-containing cell suspensions (Fig. 3 B and C), indicating brevicidine does not disrupt the membrane integrity at concentrations of 2×MIC, 1×MIC, and 0.5×MIC.

Although brevicidine showed no effect on the bacterial membrane integrity, our previous study had shown that brevicidine dissipated the proton motive force of bacteria as some novel antimicrobials do (14, 23, 24). The proton motive force of bacteria is essential for the generation of ATP, which is an essential bioactive compound for live bacteria (25, 26). If the proton motive force of bacteria is dissipated, the process of ATP biosynthesis will be inhibited or even terminated. To confirm that brevicidine can dissipate the proton motive force of bacteria, the ATP levels of brevicidne, at concentrations of 2×MIC, 1×MIC, and 0.5×MIC, treated bacteria were measured by using the BacTiter-Glo Microbial Cell Viability Assay (Promega) kit. The results showed that brevicidine decreased the ATP level of bacteria as expected (Fig. 3D). These results confirmed that brevicidine could dissipate the proton motive force of bacteria, which is one of the mechanisms by which brevicidine acts as a bactericidal antimicrobial.

The dissipation of the proton motive force of bacteria can result in an increase in the NADH levels of cells. Indeed, our previous study showed that the dehydrogenation of NADH can be inhibited by the addition of brevicidine (14). Considering the importance of the dehydrogenation of NADH to the electron transport chain of bacteria, we hypothesized that brevicidine treatment could result in the accumulation of ROS and thereafter kills bacteria. Therefore, the ROS levels of bacteria treated with different concentrations of brevicidine were measured by using a commercial kit (Beyotime, catalogue no. S0033S). The results show that brevicidine increases the ROS level of bacteria (Fig. 3E), which is one of the reasons that brevicidine shows bactericidal activity. This was further demonstrated by growth curve assay, which showed that the bactericidal activity of brevicidine was attenuated by the addition of the antioxidant N-Acetyl-L-cysteine (NAC, 6 mM) (Fig. 3F). Taking together, these results demonstrate that brevicidine exerts its bactericidal activity via dissipating the proton motive force of bacteria and thereafter inhibits ATP biosynthesis, inhibits the dehydrogenation of NADH, accumulates ROS in bacteria, and results in bacteria death.

To investigate the effect of brevicidine on the cell membrane in intuitive ways, we performed fluorescence microscopy, SEM, and TEM assays. The fluorescence microscopy assay results are consistent with the PI and NPN assays, which demonstrate that brevicidine is not an antimicrobial that kills pathogens by disrupting bacteria membrane integrity (Fig. 4A). Interestingly, after treatment with brevicidine (1×MIC) for 15 min, the bacterial surface fold was observed (Fig. 4B). In addition, after treatment with brevicidine (1×MIC) for 120 min, most of the bacteria were broken, evidenced by SEM and TEM images (Fig. 4 B and C). These results are consistent with the results of time killing assay (Fig. 3A), which showed that brevicidine started to kill bacteria after 30min treatments.

**Fig. 4.**
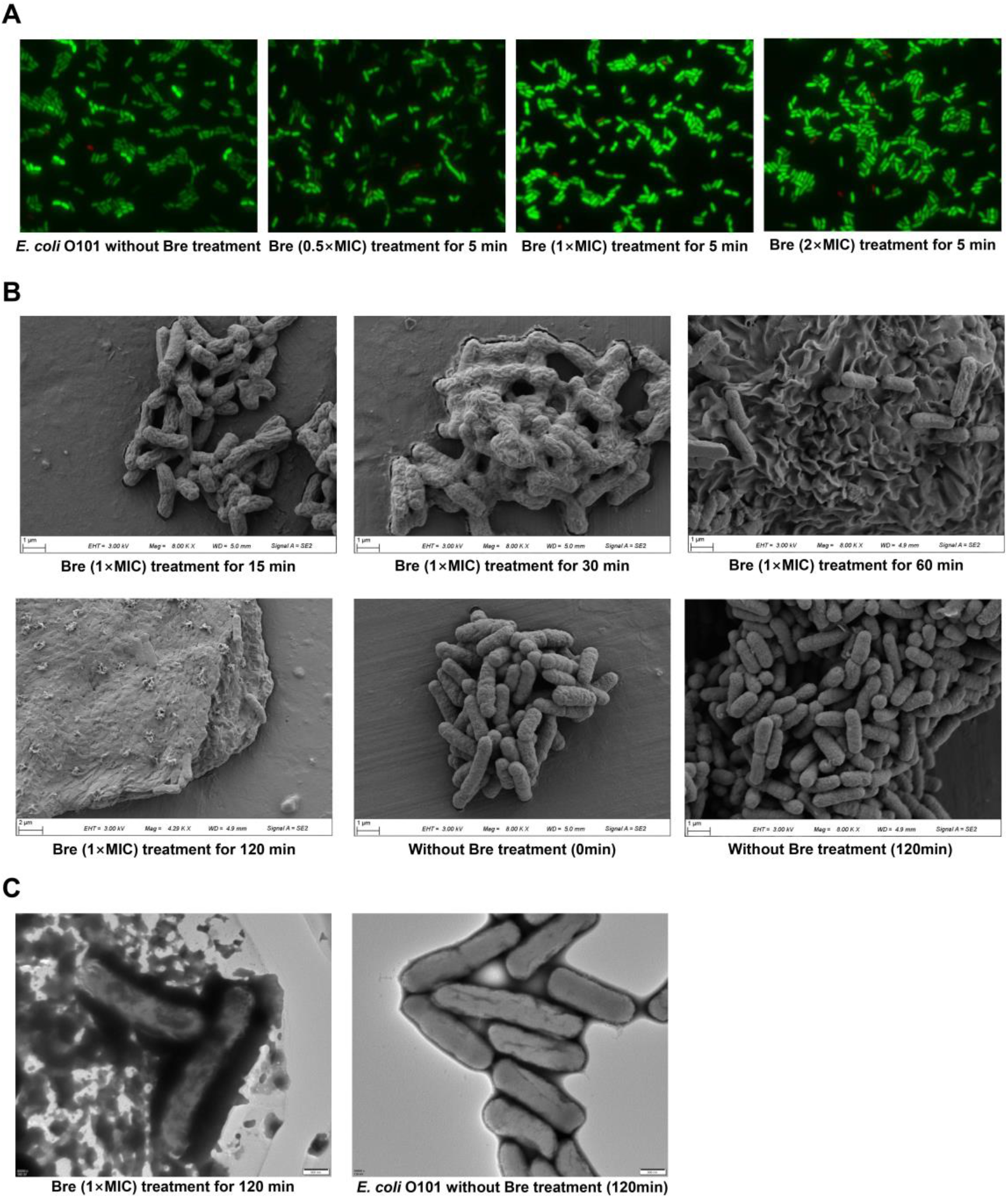
The morphology analysis of *E. coli* O101 under the treatment of brevicidine. **A**, Fluorescence microscopy images of *E. coli* O101 cells, which were challenged with brevicidine at concentrations of 2× MIC, 1× MIC, and 0.5× MIC for 5min. Green denotes a cell with an intact membrane, whereas red denotes a cell with a compromised membrane. **B**, SEM image of *E. coli* O101 cells after treatment with brevicidine at a concentration of 1× MIC for 15min, 30min, 60min, and 120min. **C**, TEM image of *E. coli* O101 cells after treatment with brevicidine at a concentration of 1× MIC for 120min.

### Brevicidine targets the membrane components LPS, PG, and CL

As brevicidine dissipates the proton motive force of bacteria, we hypothesized that brevicidine could target the component of the bacterial inner membrane. In addition, previous studies showed that brevicidine could interact with LPS (27, 28). To get an insight into the potential target, an agar diffusion assay was performed to investigate if there is an interaction between brevicidine and LPS (outer membrane component), phosphatidylethanolamine (PE, outer and inner membrane component), PG (inner membrane component), or CL (inner membrane component) (29, 30). The main component of mammalian cellular membrane phosphatidylcholine (PC) was used as a control. The results show that the addition of PC or PE did not attenuate the antimicrobial activity of brevicidine (Fig. 5A). In contrast, the antimicrobial activity was completely abolished by the addition of LPS, PG, or CL (Fig. 5A).

**Fig. 5.**
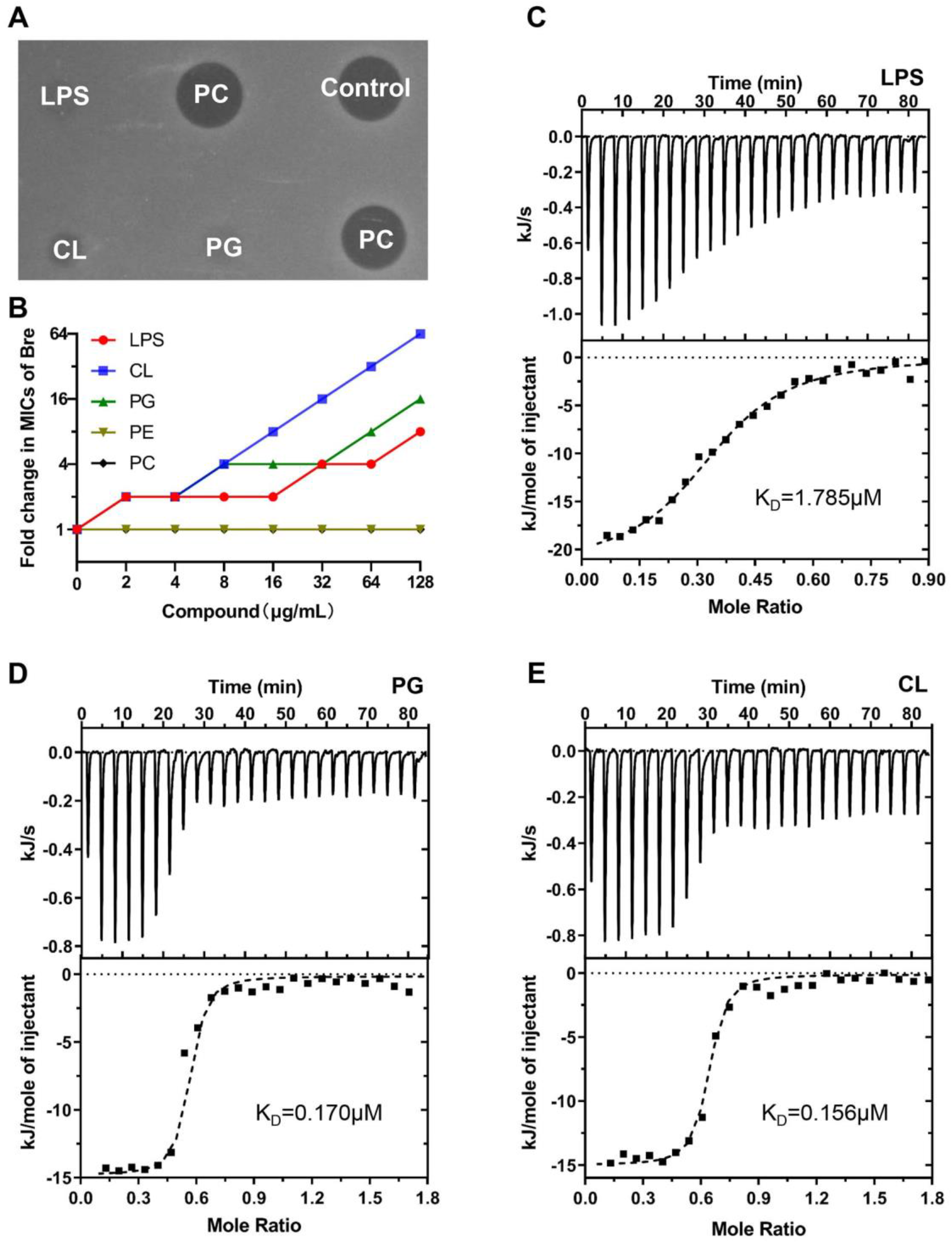
Brevicidine targets the membrane components LPS, PG, and CL. **A**, Exogenous addition of LPS, PG, or CL abolishes the antibacterial activity of brevicidine against *E. coli* O101. Inhibition zones of the mixtures of 1 nmol of brevicidine and 4 mmol of PC, LPS, PE, PG and CL, against *E. coli* O101 for overnight incubation at 37 °C. Representative examples from three technical replicates are shown. **B**, Increased MICs of brevicidine against *E. coli* O101 in the presence of LPS, PG and CL, ranging from 0 μg/ml to 128 μg/ml, based on MIC assays. Representative examples from three technical replicates are shown. **C**, ITC analysis of the interaction between brevicidine and LPS. **D**, ITC analysis of the interaction between brevicidine and PG. **E**, ITC analysis of the interaction between brevicidine and CL. The K_D_ values of brevicidine to LPS, PG and CL were calculated using the Nano Analyze Software (Waters LLC).

Subsequently, to investigate the dose-dependent effect of LPS, PG, or CL on the antimicrobial activity of brevicidine, a MIC assay was performed with different concentrations of LPS, PG, CL, PC, or PE. The MIC value of brevicidine had no changes with the addition of PC and PE, suggesting there is no interaction between brevicidine and PE (or PC) (Fig. 5B). This could explain why brevicidine targets the Gram-negative bacteria but has low cytotoxicity to mammalian cells (11, 14). LPS, PG, and CL decreased the antimicrobial activity of brevicidine in a dose-dependent manner, and the MIC value of brevicidine increased 8-fold, 16-fold, and 64-fold by the addition of 128μg/ml of LPS, PG, and CL, respectively (Fig. 5B). These results indicate that brevicidine could bind to the cell membrane components LPS, PG, and CL.

To gain an insight into the interaction between brevicidine and LPS, PG, or CL, an ITC measurement was performed. The results show that CL and PG showed comparable binding affinity to brevicidine, with K_D_ values of 0.156 μM and 0.170 Μm (Fig. 5 C and D), respectively. Notably, CL and PG have much stronger (approximately 10-fold stronger) binding affinity to brevicidine than LPS, which has a K_D_ value of 1.785 μM (Fig. 5E). These results demonstrate that brevicidine has multiple targets on the bacteria, which is good for inhibiting the development of bacterial antibiotic resistance. Together, the observed results demonstrate that brevicidine exerts its bactericidal activity via interacting with LPS in the outer membrane and targeting CL and PG in the inner membrane, and thereafter dissipating the proton motive force of bacteria.

### Brevicidine inhibits the oxidative phosphorylation and protein synthesis processes of *E. coli*

To gain an insight into the antimicrobial mechanism of brevicidine, transcriptome analysis was performed on *E. coli* treated with or without brevicidine at a concentration of 0.5μM (0.8 mg/L). The whole distribution of differentially expressed genes (DEGs) is shown in Fig. 6A. When padj value<0.05 and log2(FoldChange)>0, among 4622 genes, 587 were upregulated and 622 were downregulated, and 3413 genes were not significantly changed (Fig. 6A). Subsequently, Kyoto Encyclopedia of Genes and Genomes (KEGG) enrichment analysis was performed on the genes that had significantly expression changes. The results showed that ribosome, oxidative phosphorylation, and aminoacyl-tRNA biosynthesis pathways were significantly (padj value<0.05) affected by brevicidine treatment (Fig. 6B). After *E. coli* exposure to brevicidine for 1h oxidative phosphorylation related genes including ATP synthase, NADH-quinone oxidoreductase, and cytochrome oxidase were significantly downregulated (Fig. 6C). This change would contribute to the lower level of ATP, higher level of NADH and ROS of the brevicidine treated cells. The downregulated tRNA ligase and ribosome synthesis metabolism indicate that the protein synthesis process was inhibited by brevicidine treatment.

**Fig. 6.**
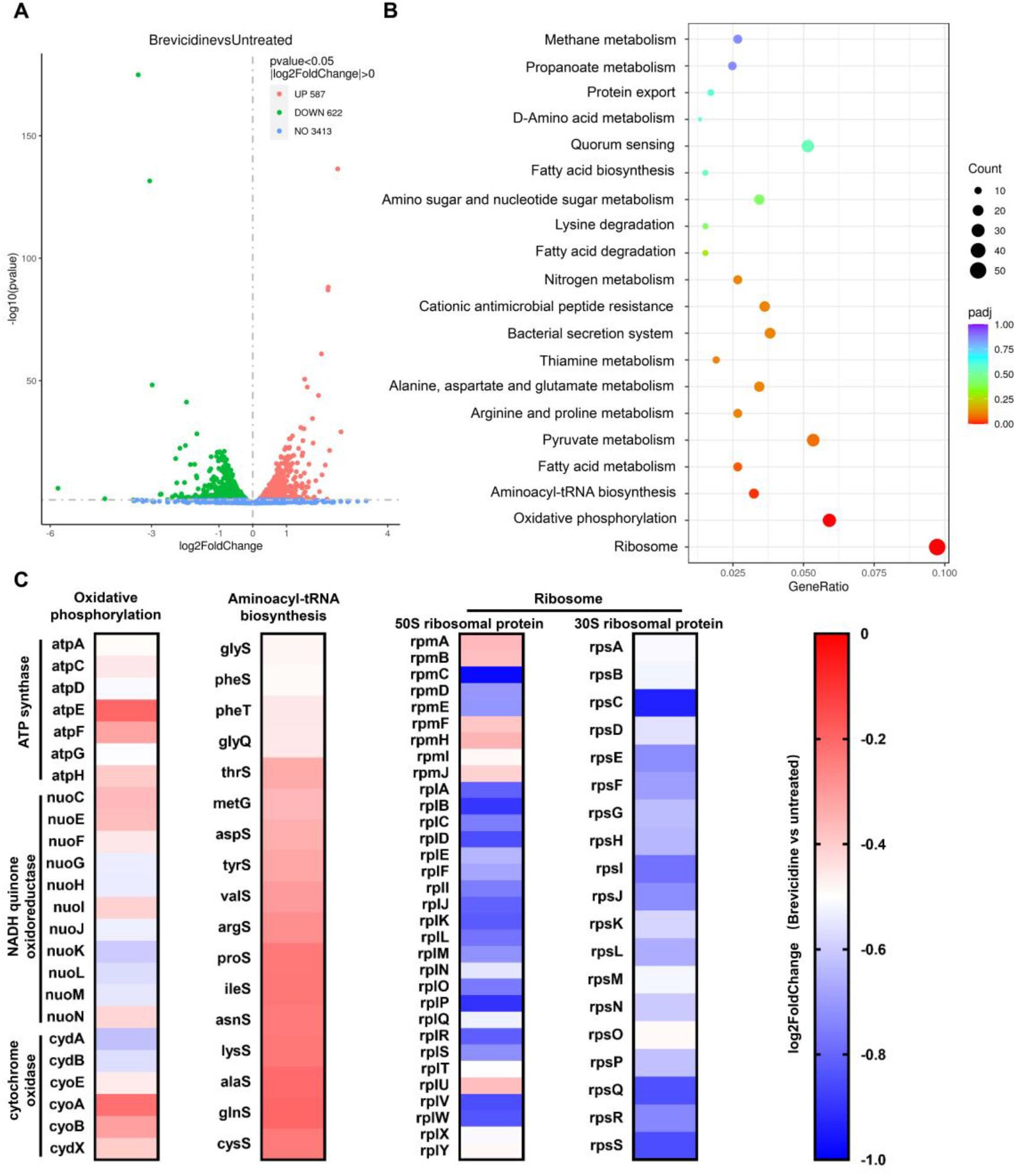
Transcriptome analysis of *E. coli* O101 under the treatment of brevicidine [0.5 μM (0.8 mg/L)] for 1 h. **A**, The volcano map of DEGs. The abscissa represents the multiple changes of gene expression between brevicidine treated and untreated *E. coli* O101; the ordinate represents the statistical significance of the change of gene expression (pvalue<0.05 |log2FoldChange|>0). **B**, Scatter plot of KEGG pathway enrichment analysis of DEGs (Brevicidine vs untreated). The vertical axis represents the KEGG enrichment analysis pathway name, and the horizontal axis represents the gene ratio of the genes in the pathway. The size of the dots indicates the number of DEGs in the pathway, and the color of the dots corresponds to different padj value ranges. **C**, Genes involved in oxidative phosphorylation, ribosome, and aminoacyl-tRNA biosynthesis were identified as significantly different between the treatment of brevicidine and untreated, with pvalue values <0.05 and log2FoldChange <0 in expression level. All genes that had significantly changes were down-regulated in oxidative phosphorylation, ribosome, and aminoacyl-tRNA biosynthesis pathways. All data are presented as means (n= 3 biological replicates).

Collectively, our findings demonstrate that brevicidine exerts its bactericidal activity via interacting with LPS in the outer membrane and targeting CL and PG in the inner membrane, and thereafter dissipation of the proton motive force of bacteria. This can result in metabolic perturbation, including inhibits ATP biosynthesis, inhibits the dehydrogenation of NADH, accumulates ROS in bacteria, and inhibits protein synthesis (Fig. 7).

**Fig. 7.**
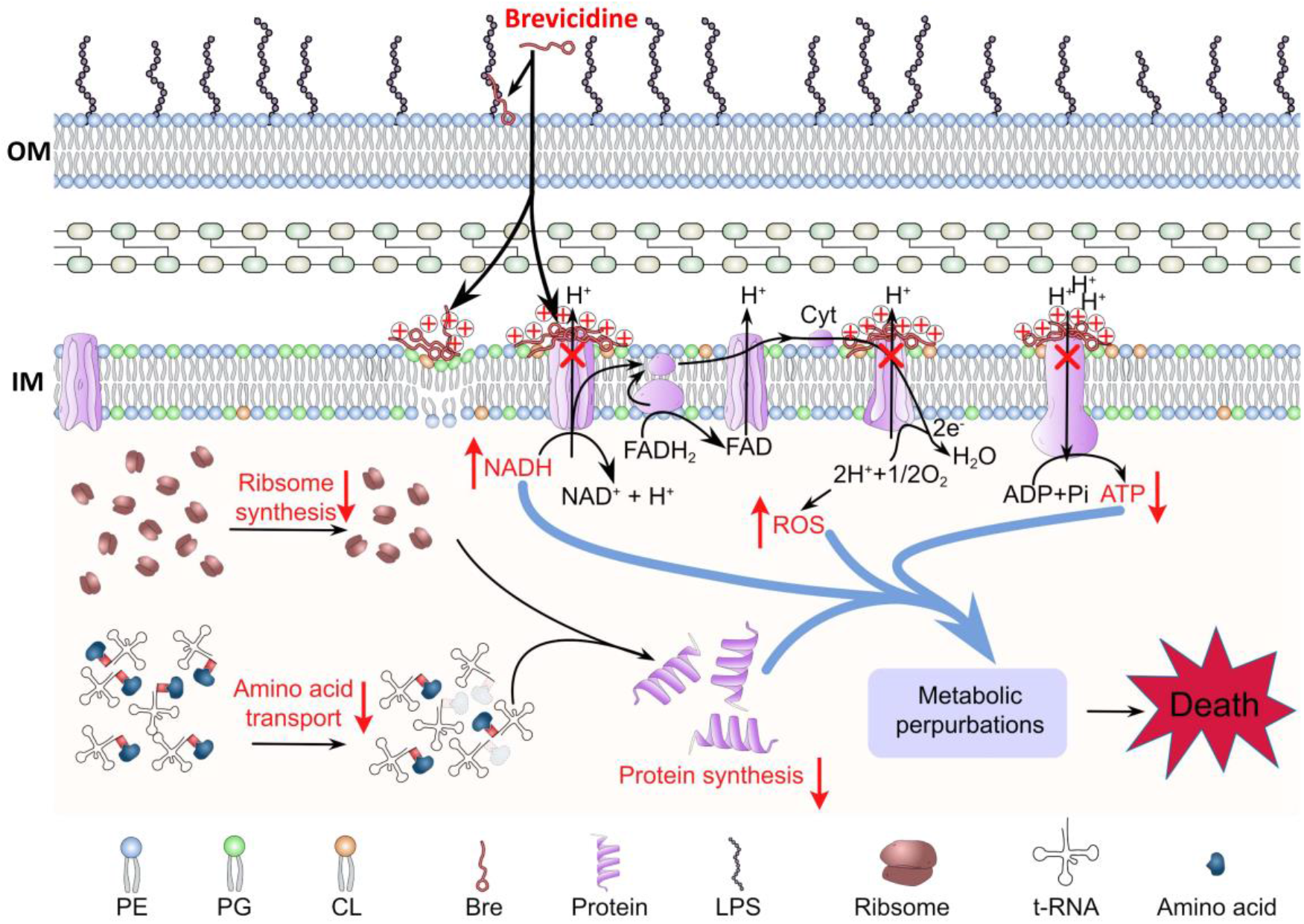
Scheme of the proposed mechanism of action brevicidine against *E. coli*. Brevicidine exerts its bactericidal activity via interacting with LPS in the outer membrane (OM) and targeting CL and PG in the inner membrane (IM), which could make a positive potential surround the outside surface of inner membrane (Note: no evidence shows brevicidine directly interacts with these channel proteins), and thereafter dissipating the proton motive force of the bacteria. This results in metabolic perturbation, including decreases in the intracellular ATP level, inhibits the dehydrogenation of NADH and accumulates ROS in bacteria. In addition, transcriptome analysis results showed that brevicidine inhibits the synthesis of t-RNA ligase (essential for amino acid transport) and ribosome protein synthesis, which can result in protein synthesis inhibition. All these actions together lead to the death of Gram-negative pathogens.

### Brevicidine showed a good therapeutic effect in a mouse peritonitis–sepsis model

Given the attractive mode of action and potent antimicrobial and anti-biofilm activity of brevicidine, we investigated its potential as a therapeutic in a mouse peritonitis–sepsis model (Fig. 8A). Mice were infected intraperitoneally with *E. coli* at a dose that leads to 80% of death. At 1h post-infection, brevicidine was introduced at single intravenous doses ranging from 10mg/kg to 40mg/kg. All brevicidine-treated mice were survived (Fig. 8B). Consistently, the bacterial load in different organs of mice significantly reduced under brevicidine treatment in a dose-dependent manner at either 1 or 7 days post-infection (Fig. 8 C-G, and Supplementary Fig. S1). These results are consistent with the previous study, which showed that brevicidine had good antimicrobial activity in a mouse thigh model (11). Together, these findings demonstrate the potential of brevicidine as a novel therapeutic antimicrobial in AMR Gram-negative pathogen infections, in particular, AMR *Enterobacteriaceae* pathogen infections.

**Fig. 8.**
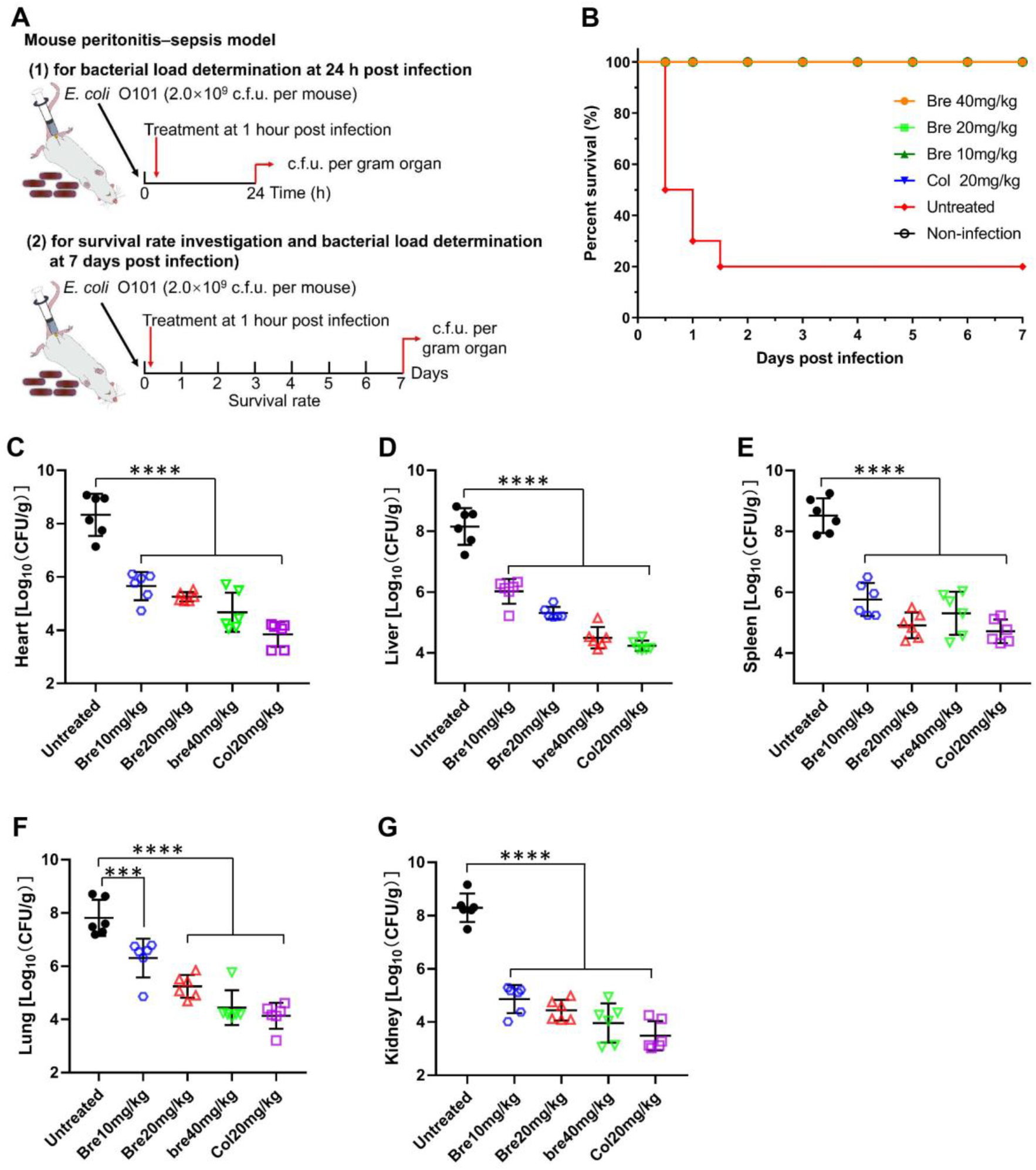
Brevicidine showed a good therapeutic effect in a mouse peritonitis–sepsis model. **A**, Scheme of the experimental protocol for the mouse peritonitis–sepsis model. **B**, Survival rates of mice in the mouse peritonitis–sepsis model (n= 10). Increased survival rates of mice over 7 d by a dose that leads to 80% of death of *E. coli* O101 (2.0 × 10^9^ c.f.u.), treated with brevicidine (10 mg per kg, 20 mg per kg, or 40 mg per kg) or with colistin (20 mg per kg). **C to G**, Brevicidine significantly reduced the bacterial load of organs of the peritonitis-sepsis mouse. At 24h post-infection, the survived mice (n = 6) were euthanized by cervical dislocation. Bacterial loads (Log10 CFU per gram of *E. coli* O101) of the heart (**C**), liver (**D**), spleen (**E**), lung (**F**) and kidney (**G**) were counted. All data were presented as means ± Standard Deviation (n = 6). Correlation analyses were evaluated by Pearson r2, ns: ***p < 0.01 and ****p < 0.001 vs. untreated group.

## Conclusions

AMR poses a major threat to human health around the world; many bacterial infections have become increasingly difficult, or even impossible, to treat with conventional antimicrobials. Therefore, new first-in-class antibiotics, operating via novel modes of action, are desperately-needed. Here, we show that brevicidine has potent antimicrobial activity against AMR *Enterobacteriaceae* pathogens, with a MIC value range of 0.5μM (0.8mg/L) to 2μM (3.0mg/L). In addition, brevicidine showed potent anti-biofilm activity against the *Enterobacteriaceae* pathogens, with the same 100% inhibition and eradication concentration of 4μM (6.1mg/L). Mechanistic studies showed that brevicidine exerts its potent bactericidal activity via interacting with LPS in the outer membrane, targeting PG and CL in the inner membrane, and dissipating the proton motive force of bacteria. This results in metabolic perturbation, including inhibition of ATP biosynthesis, inhibition of the dehydrogenation of NADH, accumulation of ROS in bacteria, and inhibition of protein synthesis (Fig. 7). Lastly, brevicidine showed a good therapeutic effect in a mouse peritonitis–sepsis model. Our findings pave the way for further research on clinical applications of brevicidine, to combat the prevalent infections caused by AMR Gram-negative pathogens worldwide.

## Materials and Methods

### Ethical approval

The animal experiment protocol was approved by the Animal Ethical and welfare Committee of Sichuan Agricultural University [permission number 20220087].

### Bacterial strains used and growth conditions

Bacterial strains used in this study are listed in S1 Table. All bacterial strains were inoculated in LB and incubated at 37°C with aeration at 220 rpm for preparing the overnight cultures.

### Purification of brevicidine

Methods for the purification of brevicidine have been described in detail previously (12). The matrix-assisted laser desorption ionization-time-of-flight mass spectrometry data and high-performance liquid chromatograph trace of purified brevicidine are shown in Supplementary Figs. S2 and S3, which are showing the correct molecular weight and high purity of the purified brevicidine.

### Minimum inhibitory concentration (MIC) assay

MIC values of antimicrobials against the tested bacterial pathogens were determined according to the standard guidelines (31). Briefly, cell concentration was adjusted to approximately 5×10^5^ cells per ml in cation-adjusted Mueller-Hinton broth (MHB). Subsequently, antimicrobials were added to the above bacterial cultures at different concentrations. After 24h of incubation at 37°C, the MIC value was defined as the lowest concentration of antimicrobial with no visible growth.

For measuring the MIC values of antimicrobials against indicator strains in the presence of cell membrane components, MHB was replaced with MHB containing different concentrations of cell membrane components, and the effect of these two conditions was compared.

### Biofilm inhibition assay

Biofilm inhibition assay was performed according to a previously described protocol (19). Briefly, an overnight culture of *E. coli* O101 was diluted 100-fold in tryptone soy broth (TSB) and incubated at 37 °C with aeration at 220 r.p.m. Bacteria were grown to an OD_600_ of 0.8, and then the concentration of cells was adjusted to an OD_600_ of 0.04 in TSB. The cell suspension was added to a 96-well plate, and antimicrobials were added at different final concentrations. After incubation at 37 °C for 24 h, the wells were washed with MiliQ water three times, and the adhered biomasses were stained with 105 μl of a 0.1% crystal violet (CV) (wt/vol) solution. The plate was placed on a low-speed orbital shaker and allowed to stain at room temperature for 30 min. Subsequently, the plate was washed with MiliQ water three times, and then 70% (vol/vol) ethanol was added to the wells (110 μl per well) and incubated for 30 min at room temperature with gentle shaking. The absorbance was measured by using a Thermo Scientific Varioskan™ LUX multimode microplate reader with a wavelength of 595 nm. The inhibition rates were calculated by the raw CV absorbance values from each growth condition vs the raw CV absorbance value of normally grown biofilm. Representative examples from three technical replicates are shown.

### Biofilm eradication assay

Biofilm eradication assay was performed according to a previously described protocol (19). Briefly, an overnight culture of *E. coli* O101 was diluted 100-fold in tryptone soy broth (TSB) and incubated at 37 °C with aeration at 220 r.p.m. Bacteria were grown to an OD_600_ of 0.8, and then the concentration of cells was adjusted to an OD_600_ of 0.04 in TSB. The cell suspension was added to 96-well plates and incubated at 37 °C for 24 h. After washing the wells with TSB three times, antimicrobials were added to the wells at different concentrations, and the plates (one for CV staining, the other one for TTC staining) were incubated at 37 °C for 24 h. For measuring the metabolic activity of biofilm, a final concentration of 0.05% TTC was added to wells with antimicrobials. Subsequently, the adhered biomasses were measured as the method described in the biofilm inhibition assay. After washing all the wells of the TTC plate three times with distilled water, dimethyl sulfoxide (DMSO) was added to the wells at 205 μl per well. After that, the plate was placed on an orbital shaker rotating at low speed and mixed at room temperature for 30 min. The absorbance of the TTC plate was measured by using a Thermo Scientific Varioskan™ LUX multimode microplate reader with a wavelength of 500 nm. The biofilm eradication (Biomass and metabolism) rate was calculated by the method described in the previous study (19). Representative examples from three technical replicates are shown.

### Time-dependent killing assay

The time-dependent killing assay was performed according to the procedure described previously (20, 21). An overnight culture of *E. coli* O101 was diluted 100-fold in MHB and incubated at 37 °C with aeration at 220 r.p.m. Bacteria were grown to an OD of 0.6, and then the concentration of cells was adjusted to 7.6×10^7^ cells per mL. Bacteria were then challenged with antimicrobials at concentrations of 1×MIC, 2×MIC, or 4×MIC in glass culture tubes at 37 °C with aeration at 220 r.p.m. Bacteria not treated with antimicrobials were used as normal growth control. At desired time points, two hundred μl aliquots were taken, centrifuged at 5,000 g for 5 min, and resuspended in 200 μl of MHB. Ten-fold serially diluted samples were plated on MHA plates. After incubation at 37 °C for 24 h, colonies were counted, and the colony forming units (c.f.u.) per ml were calculated. Each experiment was performed in triplicate.

### Membrane integrity assay

A fresh culture of *E. coli* O101 was pelleted at 5,000 g for 5 min and washed three times with MHB. The cell density was normalized to an OD_600_ of 0.2, loaded with propidium iodide (final concentration 2.5 μg per ml), and incubated for 5 min in the dark for probe fluorescence to stabilize. After the cell suspension was added to a 96-well microplate, brevicidine was added to final concentrations of 2×MIC, 1×MIC, or 0.5×MIC, with the antimicrobials added after ∼30 s, and fluorescence was monitored for 15 min. The excitation and emission wavelengths on the fluorescence spectrometer were adjusted to 533 nm and 617 nm, respectively. Representative examples from three technical replicates are shown.

### Outer membrane permeability assay

The integrity of the outer membrane was investigated with the fluorescent probe N-Phenyl-1-naphthylamine (NPN, Aladdin). A fresh culture of *E. coli* O101 was pelleted at 5,000 g for 5 min and washed three times with 10 mM HEPES containing 10 mM glucose (GHEPES, pH 7.2). The cell density was normalized to an OD_600_ of 0.2 in GHEPES, loaded with NPN (final concentration 30 μM), and incubated for 30 min in the dark for probe fluorescence to stabilize. After the cell suspension (190 μL) was added to a 96-well microplate, brevicidine (10 μL) was added to final concentrations of 2×MIC, 1×MIC, or 0.5×MIC, with the antimicrobials added after ∼30 s, and fluorescence was monitored for 15 min. The excitation and emission wavelengths on the fluorescence spectrometer were adjusted to 350 nm and 420 nm, respectively. Representative examples from three technical replicates are shown.

### Fluorescence microscopy assay

This assay was performed according to the procedure described previously (32, 33). *E. coli* O101 was grown to an OD_600_ of 0.6 in MHB. The culture was pelleted at 5,000 g for 5 min and washed with MHB three times. After normalization of the cell density to an OD_600_ of 0.2 in MHB, brevicidine was added to a final concentration of 2×MIC, 1×MIC or 0.5×MIC. After incubation at 37 °C for 5min, cells were collected by centrifugation. Subsequently, SYTO® 9 and propidium iodide (LIVE/DEAD Baclight Bacterial Viability Kit, Invitrogen) were added to the above cells. After incubation at room temperature for 15 min, cells were washed three times with MHB. Then the cell suspensions were loaded on 1.5 % agarose pads and analyzed by Nikon 80i (Japan) microscope.

### ATP determination

The ATP levels were determined using a BacTiter-Glo Microbial Cell Viability Assay kit (Promega). A fresh culture of *E. coli* O101 was pelleted at 5,000 g for 5 min and washed three times with MHB. The cell density was normalized to an OD_600_ of 0.2, loaded with brevicidine at final concentrations of 2×MIC, 1×MIC, or 0.5×MIC. At desired time points, 100 μl of cell culture were taken out, mixed with 100 μl BacTiter-Glo reagent, and incubated for 5 minutes at room temperature. Luminescence was measured with an Infinite M2000pro Microplate reader (Tecan). CCCP (40 μg/mL) was used as a positive control. The relative ATP levels were calculated using the measured Luminescence values vs the Luminescence value of untreated cells at relative time points.

### Reactive oxygen species (ROS) measurement

The levels of ROS in *E. coli* O101 treated with different concentrations of brevicidine were measured by 10 μM of 2′,7′-dichlorofluorescein diacetate (DCFH-DA), following the manufacturer’s instruction (Beyotime, catalogue no. S0033S). Briefly, a fresh culture of *E. coli* O101 was pelleted at 5,000 g for 5 min and washed three times with MHB. The cell density was normalized to an OD_600_ of 1.0, loaded with DCFH-DA at a final concentration of 10 μM, and the mixture incubated at 37 °C for 30 min. After washing with MHB three times, 190 μl of probe-labeled bacterial cells were added to a 96-well plate, and then 10 μl of brevicidine. Fluorescence was recorded by using a Thermo Scientific Varioskan™ LUX multimode microplate reader with the excitation wavelength at 488 nm and the emission wavelength at 525 nm. The antioxidant N-Acetyl-L-cysteine (NAC, 6 mM) was used as a control to neutralize the production of ROS.

### Scanning electron microscopy (SEM) and transmission electron microscopy (TEM) assays

The morphology of 1μM (1.52 mg/L) brevicidne treated *E. coli* O101 was recorded in vacuum on a ZEISS Gemini SEM300 with an acceleration voltage of 3kV. For TEM measurements, samples were prepared by delivering 2μl of 1μM (1.52 mg/L) brevicidine treated *E. coli* O101 suspension to carbon-coated copper grids and dried in a vacuum system. The morphology was obtained under a JEM-2100Plus Electron Microscope using an acceleration voltage of 120kV.

### Spot-on-lawn assay

An overnight cultured *E. coli* O101 was added to 0.7% MHA (w/v, temperature 42 °C) at a final concentration of 0.25% (v/v), and then the mixture was poured onto the plates, with 20 mL for each 15 cm diameter circular plate. Subsequently, a spot-on-lawn assay was used to analyze the antimicrobial activity of brevicidine in the presence of cell membrane components (20). In short, 20 μL brevicidine (50 μM) containing 200 μM lipopolysaccharides (LPS, from Escherichia coli 055:B5, Solarbio, catalogue no. L8880, ≥98%), L-α-phosphatidylcholine, (PC, Aladdin, catalogue no. L130331, ≥99%), L-α-phosphatidylglycerol (PG, Aladdin, catalogue no. L130372; ≥99%), L-α-phosphatidylethanolamine (PE, Aladdin, catalogue no. L130310; ≥99%) or Cardiolipin (CL, Aladdin, catalogue no. B130227;≥99%) was loaded to the agar plate that contains indicator strain. After the brevicidiene/cell membrane component mixture solution drops had dried, the plates were transferred to a 37 °C incubator for overnight incubation.

### Isothermal Titration Calorimetry (ITC) assay

ITC was performed with the Low Volume NanoITC (TA Instruments) at 25°C to determine the interaction between brevicidine and LPS, PG, or CL. Brevicidine was diluted in a buffer (20 mM HEPES, pH7.0) to a final concentration of 50 μM. Samples were degassed before use. The chamber was filled with 300 μL of the brevicidine solution, and LPS (250μM), PG (500μM), or CL (500μM) was titrated into the chamber at a rate of 1.96 μL/200 s with a stirring rate of 300 rpm. The control experiment was performed with 20 mM HEPES titration to 50 μM brevicidine. The K_D_ values of brevicidine to LPS, PG, or CL were calculated using the Nano Analyse Software (Waters LLC).

### Transcriptome analysis of *E. coli* O101 treated with brevicidine

Three replicates of overnight bacteria cultures were diluted at 1:100 in MHB and grown at 37°C with aeration at 220rpm. Bacteria were grown to an OD_600_ of 0.4, and then the concentration of cells was adjusted to an OD_600_ of 0.2 in MHB. Subsequently, the bacteria were treated with brevicidine at a final concentration of 0.5μM (0.8 mg/L). Bacteria without brevicidine treatment were used as untreated control. Three replicates were performed for both brevicidine treated and untreated bacterial cultures, 4ml for each sample. After incubation at 37°C with aeration at 220rpm for 1h, the bacteria from different treatments were collected by centrifuging, and the bacterial pallets were stored in liquid nitrogen immediately. Subsequently, the samples were sent to Novogene Co., Ltd (China) for transcriptome analysis. After RNA extraction, mRNA purification, and cDNA synthesis, the samples were sequenced on the Illumina NovaSeq6000. The quality of the resulting fastq reads was mapped on the reference genome using Bowtie2 2.3.4.3 using default settings. Feature Counts 1.5.0-p3 was used to get the gene counts. DESeq2 1.20 was used to identify all genes, and padj ≤0.05 and |log2 (foldchange) | ≥ 0 of genes between two treatments were regarded as differentially expressed genes (DEGs). The volcano map and KEGG enrichment of DEGs were performed with Magic Novogene package (https://magic.novogene.com). The RNA-Seq data have been deposited in the NCBI Gene Expression Omnibus with the accession number PRJNA872022.

### Mouse peritonitis–sepsis model

Brevicidine was tested against *E. coli* O101 in a mouse peritonitis–sepsis model to assess its *in vivo* bioavailability. BALB/c male mice (n=6 per group) were infected with 0.4ml of bacterial suspension (2×10^9^ c.f.u. per mouse) via intraperitoneal injection, a concentration that achieves approximately 80% mortality within 48 h post-infection. At 1h post-infection, mice were treated with 0.9% NaCl, brevicidine (40mg/kg), brevicidine (20mg/kg), brevicidine (10mg/kg), or colistin (20mg/kg) via intravenous injection. Mice without *E. coli* O101 infections were used as the normal group control. Once the infected mice died, different organs, including the heart, liver, spleen, lung, and kidney, were collected and homogenized in sterilized PBS for bacterial load quantification. At 24h post-infection, the organs of the survived mice were collected to measure the bacterial load. In addition, the same mouse peritonitis–sepsis model was determined for 7d (n=10), and the bacterial load in the organs of surviving mice was quantified at 7 days post-infection.

### Statistical analysis

GraphPad Prism 8.0 was used to fit the data of Fig. 2, Fig. 3, Fig. 5, Fig. 8, and Supplementary Fig. S1. The statistical significance of the data was assessed using a two-tailed Student’s t-test with GraphPad Prism 8.0. Correlation analyses were evaluated by Pearson r2, ns: p>0.05, *p<0.05, **p<0.01, ***p<0.001, and ****p<0.0001.

## Data Availability Statement

The transcriptome (RNA sequencing) data that support the findings of this study have been deposited in the NCBI Sequence Read Archive (SRA) with the accession code PRJNA872022.

## Finding

Xinghong Zhao was supported by the Science and Technology Project of Sichuan Province (2022YFH0057). Hongping Wan was supported by the Science and Technology Project of Sichuan Province (2022YFH0062).

## Acknowledgments

We thank Prof. Kui Zhu (National Center for Veterinary Drug Safety Evaluation, College of Veterinary Medicine, China Agricultural University, Beijing, China.) for providing the multidrug resistant *Escherichia coli* (B2 and 16QD) strains.

## References

1. O’neill, J. I. M. (2014) Antimicrobial resistance: tackling a crisis for the health and wealth of nations. Rev. Antimicrob. Resist. 1, 1–16

2. Kwon, J. H., and Powderly, W. G. (2021) The post-antibiotic era is here. Science. 373, 471

3. Knight, G. M., Glover, R. E., McQuaid, C. F., Olaru, I. D., Gallandat, K., Leclerc, Q. J., Fuller, N. M., Willcocks, S. J., Hasan, R., and van Kleef, E. (2021) Antimicrobial resistance and COVID-19: Intersections and implications. Elife. 10, e64139

4. Strathdee, S. A., Davies, S. C., and Marcelin, J. R. (2020) Confronting antimicrobial resistance beyond the COVID-19 pandemic and the 2020 US election. Lancet. 396, 1050–1053

5. Lai, C.-C., Chen, S.-Y., Ko, W.-C., and Hsueh, P.-R. (2021) Increased antimicrobial resistance during the COVID-19 pandemic. Int. J. Antimicrob. Agents. 57, 106324

6. Rizvi, S. G., and Ahammad, S. Z. (2022) COVID-19 and antimicrobial resistance: A cross-study. Sci. Total Environ. 807, 150873

7. Butler, M. S., Blaskovich, M. A. T., and Cooper, M. A. (2017) Antibiotics in the clinical pipeline at the end of 2015. J. Antibiot. (Tokyo). 70, 3

8. Butler, M. S., Blaskovich, M. A., and Cooper, M. A. (2013) Antibiotics in the clinical pipeline in 2013. J. Antibiot. (Tokyo). 66, 571

9. Roemer, T., and Boone, C. (2013) Systems-level antimicrobial drug and drug synergy discovery. Nat. Chem. Biol. 9, 222

10. Süssmuth, R. D., and Mainz, A. (2017) Nonribosomal peptide synthesis—principles and prospects. Angew. Chemie Int. Ed. 56, 3770–3821

11. Li, Y.-X., Zhong, Z., Zhang, W.-P., and Qian, P.-Y. (2018) Discovery of cationic nonribosomal peptides as Gram-negative antibiotics through global genome mining. Nat. Commun. 9, 3273

12. Zhao, X., Li, Z., and Kuipers, O. P. (2020) Mimicry of a Non-ribosomally Produced Antimicrobial, Brevicidine, by Ribosomal Synthesis and Post-translational Modification. Cell Chem. Biol. 27, 1262-1271.e4

13. Al Ayed, K., Ballantine, R. D., Hoekstra, M., Bann, S. J., Wesseling, C. M. J., Bakker, A. T., Zhong, Z., Li, Y.-X., Brüchle, N. C., and van der Stelt, M. (2022) Synthetic studies with the brevicidine and laterocidine lipopeptide antibiotics including analogues with enhanced properties and in vivo efficacy. Chem. Sci. 13, 3563–3570

14. Zhao, X., and Kuipers, O. P. (2021) BrevicidineB, a new member of the brevicidine family, displays an extended target specificity. Front. Microbiol. 12, 1482

15. Tacconelli, E., Magrini, N., Kahlmeter, G., and Singh, N. (2017) Global priority list of antibiotic-resistant bacteria to guide research, discovery, and development of new antibiotics. World Heal. Organ. 27, 318–327

16. Song, M., Liu, Y., Huang, X., Ding, S., Wang, Y., Shen, J., and Zhu, K. (2020) A broad-spectrum antibiotic adjuvant reverses multidrug-resistant Gram-negative pathogens. Nat. Microbiol. 5, 1040–1050

17. Liu, Y., Shi, L., Su, L., van der Mei, H. C., Jutte, P. C., Ren, Y., and Busscher, H. J. (2019) Nanotechnology-based antimicrobials and delivery systems for biofilm-infection control. Chem. Soc. Rev. 48, 428–446

18. Sharma, G., Sharma, S., Sharma, P., Chandola, D., Dang, S., Gupta, S., and Gabrani, R. (2016) Escherichia coli biofilm: development and therapeutic strategies. J. Appl. Microbiol. 121, 309–319

19. Haney, E. F., Trimble, M. J., and Hancock, R. E. W. (2021) Microtiter plate assays to assess antibiofilm activity against bacteria. Nat. Protoc. 16, 2615–2632

20. Zhao, X., Yin, Z., Breukink, E., Moll, G. N., and Kuipers, O. P. (2020) An Engineered Double Lipid II Binding Motifs-Containing Activity against Enterococcus faecium. Antimicrob. Agents Chemother. 64, 1–12

21. Ling, L. L., Schneider, T., Peoples, A. J., Spoering, A. L., Engels, I., Conlon, B. P., Mueller, A., Schäberle, T. F., Hughes, D. E., and Epstein, S. (2015) A new antibiotic kills pathogens without detectable resistance. Nature. 517, 455

22. Helander, I. M., and Mattila‐Sandholm, T. (2000) Fluorometric assessment of Gram‐negative bacterial permeabilization. J. Appl. Microbiol. 88, 213–219

23. Cochrane, S. A., Findlay, B., Bakhtiary, A., Acedo, J. Z., Rodriguez-Lopez, E. M., Mercier, P., and Vederas, J. C. (2016) Antimicrobial lipopeptide tridecaptin A1 selectively binds to Gram-negative lipid II. Proc. Natl. Acad. Sci. 113, 11561–11566

24. Stokes, J. M., Yang, K., Swanson, K., Jin, W., Cubillos-Ruiz, A., Donghia, N. M., MacNair, C. R., French, S., Carfrae, L. A., and Bloom-Ackerman, Z. (2020) A deep learning approach to antibiotic discovery. Cell. 180, 688–702

25. Ahmed, S., and Booth, I. R. (1983) The use of valinomycin, nigericin and trichlorocarbanilide in control of the protonmotive force in Escherichia coli cells. Biochem. J. 212, 105–112

26. Bakker, E. P., and Mangerich, W. E. (1981) Interconversion of components of the bacterial proton motive force by electrogenic potassium transport. J. Bacteriol. 147, 820–826

27. Malina, A., and Shai, Y. (2005) Conjugation of fatty acids with different lengths modulates the antibacterial and antifungal activity of a cationic biologically inactive peptide. Biochem. J. 390, 695–702

28. Li, Y. X., Zhong, Z., Zhang, W. P., and Qian, P. Y. (2018) Discovery of cationic nonribosomal peptides as Gram-negative antibiotics through global genome mining. Nat. Commun. 9, 2–10

29. Sohlenkamp, C., and Geiger, O. (2016) Bacterial membrane lipids: diversity in structures and pathways. FEMS Microbiol. Rev. 40, 133–159

30. Santos, R. S., Figueiredo, C., Azevedo, N. F., Braeckmans, K., and De Smedt, S. C. (2018) Nanomaterials and molecular transporters to overcome the bacterial envelope barrier: Towards advanced delivery of antibiotics. Adv. Drug Deliv. Rev. 136, 28–48

31. Wiegand, I., Hilpert, K., and Hancock, R. E. W. (2008) Agar and broth dilution methods to determine the minimal inhibitory concentration (MIC) of antimicrobial substances. Nat. Protoc. 3, 163

32. Li, Z., Chakraborty, P., de Vries, R. H., Song, C., Zhao, X., Roelfes, G., Scheffers, D., and Kuipers, O. P. (2020) Characterization of two relacidines belonging to a novel class of circular lipopeptides that act against Gram‐negative bacterial pathogens. Environ. Microbiol. 22, 5125–5136

33. Li, Z., de Vries, R. H., Chakraborty, P., Song, C., Zhao, X., Scheffers, D.-J., Roelfes, G., and Kuipers, O. P. (2020) Novel modifications of nonribosomal peptides from Brevibacillus laterosporus MG64 and investigation of their mode of action. Appl. Environ. Microbiol.

